# When your heart isn’t in it anymore: Cardiac correlates of task disengagement

**DOI:** 10.1101/2024.06.21.599851

**Authors:** Andrew W. Corcoran, Arthur Le Coz, Jakob Hohwy, Thomas Andrillon

**Affiliations:** Monash Centre for Consciousness & Contemplative Studies, Monash University, Melbourne, Victoria, VIC 3800, Australia; Sorbonne Université, Institut du Cerveau - Paris Brain Institute - ICM, Inserm, CNRS, 75013 Paris, France

**Keywords:** Arousal, Attention, Brain-body interaction, Heartbeat-evoked potential, Heart-rate variability, Mind-blanking, Mind-wandering, Pupillometry, SART, Vigilance

## Abstract

Neuroscience is beginning to uncover the role of interoceptive feedback in perception, learning, and decision-making; however, the relation between spontaneous visceral and cognitive dynamics has received surprisingly little scrutiny. Here, we investigate how subjective, physiological, and behavioural indicators of arousal and attentional state vary in relation to ongoing cardiac activity and brain-heart coupling. Electroencephalogram, electrocardiogram, and pupillometric records were obtained from 65 adults during the performance of a sustained attention to response task (SART). Thought probes were intermittently administered during the SART to collect subjective reports of attentional state (i.e., on-task, mind-wandering, mind-blanking) and vigilance level (i.e., alertness vs. sleepiness). Mind-wandering and mind-blanking reports increased in frequency with time-on-task and were accompanied by decreases in alertness and pupil-linked arousal, but evinced distinct psychophysiological and behavioural profiles. While mind-wandering was associated with greater heart-rate variability and late modulation of the heartbeat-evoked potential, mind-blanking was characterised by greater decreases in heart-rate, pupil size, and brain-heart coupling. Lower heart-rate predicted decreased vigilance and pupil size, in addition to slower, less-biased responses; increased heart-rate variability predicted more impulsive behaviour and pupil dilation. Together, these findings reveal that cardiac and brain-heart connectivity measures afford complementary information about arousal states and attentional dynamics during task performance.

## Introduction

Neural monitoring of internal bodily states (interoception) is essential for homeostasis, and conditions the way we perceive and act in the world ^1–5^. For example, visceral afferent signals generated by the heartbeat modulate the sensation ^6–11^, learning, and memory of external stimuli ^12–14^, coincide with the initiation of voluntary actions ^15–17^, and are adaptively regulated to optimise perceptual decision-making ^18^. Moreover, cortical processing of cardiac-related interoceptive signals predicts stimulus detection ^19–21^, interoceptive attention ^22–24^, and decision-making ^25^ in healthy individuals, and may harbour diagnostic information in disorders of consciousness ^26,27^. While this growing body of work demonstrates the involvement of cardio-afferent signals in shaping conscious experience and behaviour, very little is known about the way cardiac activity relates to ongoing cognitive dynamics, such as the spontaneous vacillation between task-focused attention and task disengagement.

Arousal state and attentional engagement have long been associated with cardiac regulation ^28–32^. For instance, phasic heart-rate responses to salient events vary depending on stimulus attributes such as intensity ^33,34^ and content ^35,36^, while prevailing heart-rate is reliably modulated by task demands (e.g., attentive listening vs. mental arithmetic) ^37,38^ and situational properties (e.g., uncertainty, incentivisation) ^39^. Early studies of heart-rate variability (HRV) – i.e., fluctuations in the duration separating each heartbeat ^40–42^ – also revealed sensitivity to task performance, with transient heart-rate decelerations indexing both anticipatory ^43,44^ and error-related processing ^45^ (for recent reviews, see Skora and colleagues ^5^ and Di Gregorio and colleagues ^46^). HRV has since been extensively studied as a biomarker of the capacity to regulate one’s arousal, attention, and behaviour ^47–51^, and would therefore seem to present a promising psychophysiological candidate for tracking the spontaneous evolution of cognitive dynamics.

Alterations in cardiac parameters such as heart-rate (or its reciprocal, heart period) and HRV are primarily driven by the central modulation of sympathetic and parasympathetic outflows ^52–55^. Such parameters thus afford information about the way integrated cortical and subcortical networks co-ordinate physiological and behavioural responses to evolving conditions ^56–61^. Effective regulation depends on the neural monitoring of visceral afferent feedback generated by autonomic activity ^62–64^; the heartbeat-evoked potential (HEP ^65,66^) has been widely interpreted as a neurophysiological index of such interoceptive processing ^67,68^. Intriguingly, this evoked response is modulated by attentional focus – attending to heartbeat sensations amplifies the HEP compared to when attention is directed towards external stimuli ^22–24,69–72^.

Altered heartbeat-evoked responses have also been reported in the context of self-related spontaneous cognition ^73,74^ and mental imagery ^75^, indicating that HEP amplitude is sensitive to the content of thought independent of cardiac dynamics.

A more recent development in the psychophysiology and neuroscience of brain-heart integration is the use of frequency-domain coupling measures to assess how cortical and cardiac rhythms co-evolve through time ^76^. While the HEP tracks how neural responses to the heartbeat unfold over the course of a few hundred milliseconds, coupling measures can be deployed to summarise dependencies between time-series on the order of seconds or minutes. Signal processing and modelling techniques may also be used to infer whether fluctuations in coupling strength are primarily driven by one signal or another (i.e., whether changes in brain-heart connectivity reflect a relative shift towards top-down autonomic regulation or bottom-up interoceptive processing). For example, a recent study reported a predominantly bottom-up pattern of brain-heart coupling under (task-free) resting-state conditions, whereby spontaneous neural oscillations were modulated by the phase of the HRV time-series ^77^. Brain-heart coupling has also been found to differ across levels of arousal (e.g., sleep stage ^78,79^, physiological ^80,81^ or mental challenge ^82,83^, emotion induction ^84–86^). However, we are unaware of any previous attempts to evaluate whether brain-heart coupling systematically varies as a function of attentional state dynamics.

In this study, we investigate whether cardiac parameters encode information distinguishing subjective reports of task-focused attention (‘on-task’ state) from spontaneous episodes of task disengagement. We distinguished between two modes of task disengagement – task-unrelated thought (‘mind-wandering’) and the absence of thought (‘mind-blanking’) – because of their distinct signatures in terms of behaviour and arousal, with mind-blanking being associated with lower levels of arousal and more sluggish responding than mind-wandering ^87,88^. Notably, mind-wandering has previously been linked with increased heart-rate during the performance of laboratory-based tasks ^89–92^. However, this association was not evident in a 24-hr experience sampling study ^93,94^. Short-term HRV has also been identified as a potential biomarker of mind-wandering, although differences in HRV have thus far only been reported for perseverative, negatively-valenced thoughts ^89,93,94^. We are unaware of any attempt to characterise the expression of cardiac dynamics during mind-blanking, although recent work suggests cardiac features may be informative for distinguishing mind-blanking from other states ^88^. Beyond the mind-wandering and mind-blanking literature, subjective disengagement from challenging tasks has been associated with blunted increases in heart-rate relative to resting-state ^95,96^, indicating that cardiac dynamics afford information about one’s willingness to invest effort in task performance.

Given the paucity of data on the relation between cardiac activity and spontaneous cognition (in contrast to the rich literature linking mind-wandering and pupillometry ^87,97–101^) – and extending beyond recent studies investigating brain-heart interaction under task-free resting-state conditions ^77,88,102^ and arousal state manipulations ^80–86^ – we sought to identify the cardiac signatures of task (dis)engagement during the performance of a Go/NoGo paradigm known as the sustained attention to response task (SART ^103,104^). We evaluated how cardiac parameters varied as a function of participants’ focus of attention (on-task, mind-wandering, mind-blanking) while performing the SART, and how these profiles compared with established indices of arousal and task (dis)engagement (i.e., subjective alertness/fatigue, pupil size, behavioural performance). Finally, we computed two complementary measures of interoceptive processing – the well-established HEP ^67,68^ and a more-recently proposed method for quantifying cerebro-peripheral signal coupling ^105,106^ – to assess whether attentional states are characterised by distinctive patterns of brain-heart interaction.

## Materials and Methods

### Participants

We analysed data collected from 26 adults as part of a previously published study conducted in Melbourne, Australia ^107^, and from 41 adults who participated in a replication of the original experiment in Paris, France. Both datasets featured electroencephalogram (EEG), electrocardiogram (ECG), and pupillometric recordings (except for two participants) obtained during performance of the SART. One participant from each dataset was excluded from analysis due to a faulty ECG. The remaining sample consisted of 36 males and 29 females aged 20 to 39 years (M = 27.3, S.D. = 5.3; age of one participant missing). All individuals provided written informed consent prior to participating in the study. The study protocol was approved by the Monash University Human Research Ethics Committee (Project ID: 10994) and Comité d’Éthique de la Recherche (CER) de Sorbonne Université (Project ID: CER-2023-DEGRAVE-MW_Respi).

### Stimuli and experimental design

Task instructions and stimuli were presented using the Psychophysics Toolbox (Melbourne: v3.0.14; Paris: v3.0.18 ^108^) for MATLAB (The MathWorks Inc., Natick, MA, USA; Melbourne: R2018b, Paris: 2022b). Participants viewed task stimuli while seated in a dimly-lit room with their head stabilised on a support positioned approximately 60 cm from the computer screen.

Participants performed two versions of a modified SART: one featuring face stimuli, another featuring numerical digits (Figure 1). Face stimuli were sampled from the Radboud Face Database ^109^. Eight faces (4 female) featuring a neutral expression were selected as Go stimuli; one image of a smiling female served as the NoGo stimulus. Stimuli for the Digit SART were computer-generated integers ranging from 1 to 9, with “3” serving as the NoGo stimulus. All digits were presented at a uniform font size.

**Figure 1:**
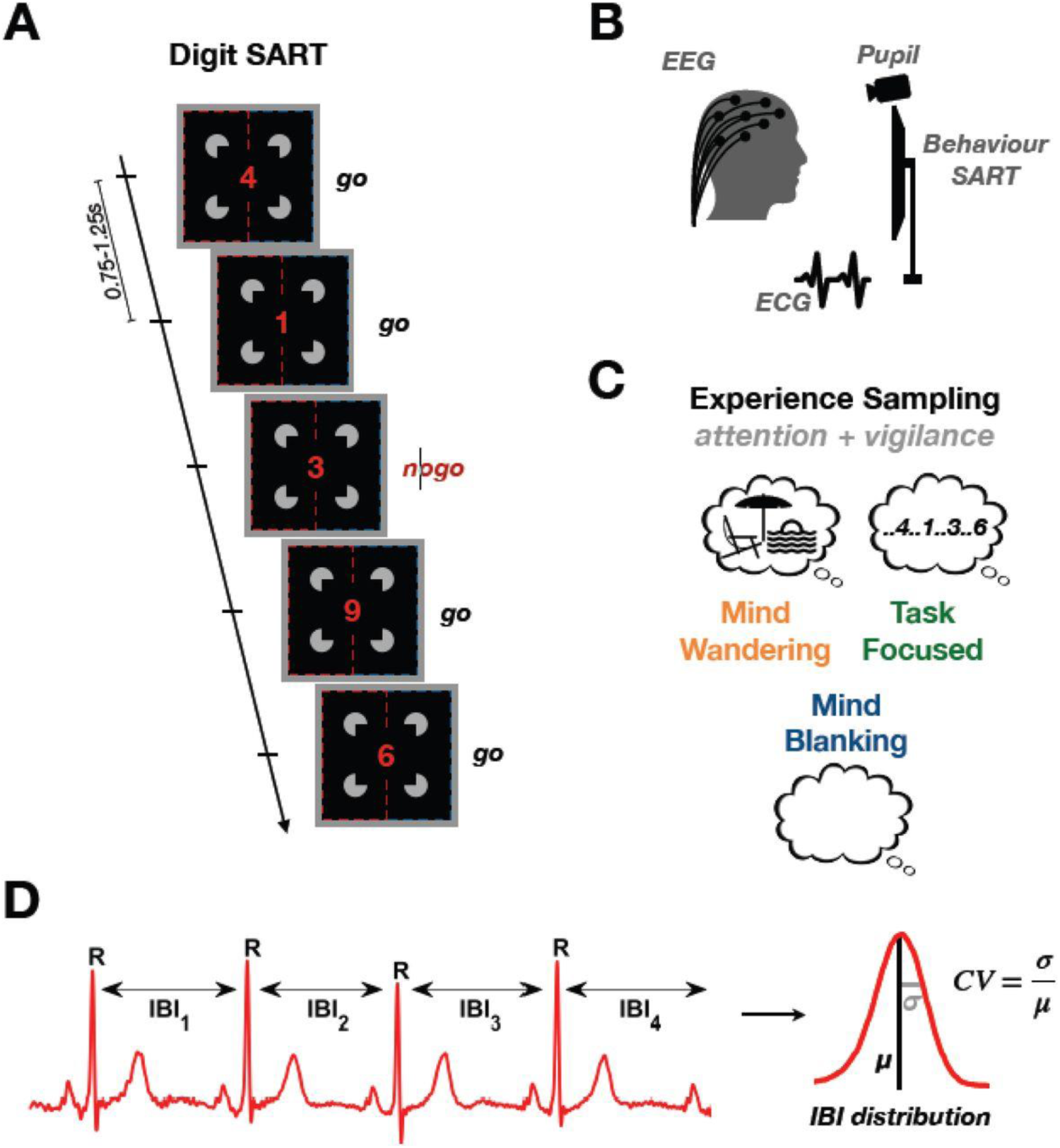
**Tracking the neural and cardiac correlates of task disengagement** A: Participants were instructed to perform a modified version of the Sustained Attention to Response Task (SART), a Go/NoGo task involving either face or digit stimuli (block design). Participants had to respond to all stimuli (Go trials) except when presented with a specific target stimulus (smiling face or digit ‘3’; NoGo trials). B: For each participant, EEG and ECG signals, pupil size, and behavioural data on the SART were continuously recorded. C: Participants were interrupted every 40-70 s (10 times per block) and asked to report their attentional state (on-task, mind-wandering, or mind-blanking) and vigilance level (alertness). D: In ECG recordings, we automatically detected R-peaks and computed the duration of successive interbeat intervals (IBIs) during SART performance. The IBI time-series was used to calculate the IBI distribution from which we derived the mean IBI duration (i.e., average heart period; inverse heart-rate) and coefficient of variation (CV; i.e., heart-rate variability) during epochs preceding thought probes.

In order to induce steady-state visual evoked potentials (SSVEPs) in the EEG, the left and right halves of each presented face stimulus were flickered at different frequencies (12 and 15 Hz; laterality counterbalanced across blocks). Digit stimuli were embedded within a Kanizsa illusory square, the left and right parts of which were likewise flickered. Analysis of this component of the experimental design is postponed to future work. Please note, flicker frequencies were sufficiently high that participants did not report any impact on their capacity to perform the SART.

Stimuli were presented continuously at the centre of the screen, with successive trial onsets occurring at intervals of 750 to 1250 ms (randomly sampled from a uniform distribution). All 9 stimuli within the stimulus set were present once in a pseudo-randomised order, before the set was randomly permuted for the next 9 trials (with the constraint that the same stimulus never appeared twice in succession). The probability of a NoGo trial was thus fixed at *p* = .11. Participants were instructed to pay attention to each presented stimulus and respond via button-press each time a Go stimulus appeared. Conversely, they were told to withhold their response whenever a NoGo stimulus was presented. Participants were advised to prioritise accuracy over speed of response, in line with evidence that this strategy improves the precision of SART errors as an index of attentional lapses ^110^.

The SART was intermittently interrupted by the presentation of the word “STOP”, followed by a set of 7 or 8 thought probes. Our analysis focuses on participants’ subjective reports of the content of their attentional focus “just before the interruption”. Participants responded to this probe by selecting one of the following options: (1) “task-focused” (i.e., on-task), (2) “off-task” (i.e., focused on something other than the SART; mind-wandering), (3) “mind-blanking” (i.e., focused on nothing), (4) “don’t remember”. In the analyses that follow, responses (3) and (4) were collapsed into a single “mind-blanking” category. Data from the final probe of each set, which asked participants to rate their level of vigilance “over the past few trials” on a 4-point Likert scale (1 = “Extremely Sleepy”; 4 = “Extremely Alert”), are also reported. In addition, participants were also asked to answer the following questions, which were not analysed here:

1. “Were you looking at the screen?” (response: yes / no); (2) “How aware were you of your focus?” (response: from 1, I was fully aware, to 4, I was not aware at all); (3) “Was your state of mind intentional?” (response: from 1, entirely intentional, to 4, entirely unintentional); (6) “How engaging were your thoughts?” (response: from 1, not engaging, to 4, very engaging); (7) “How well do you think you have been performing?” (response: from 1, not good, to 4, very good).

Prior to commencing the experiment, participants performed two blocks of practice trials (1 block of 27 trials for each variant of the SART). Performance feedback (proportion of correct responses, average response time) was provided at the end of each practice block. Participants then performed 3 blocks of each SART (block order randomly permuted per participant) for a total of 6 blocks. Blocks comprised between 459 and 616 trials (median = 540), lasted approximately 12–15 min, and were separated by self-paced breaks (total duration: M = 95.1 min, S.D. = 10.4). Each block comprised 10 sets of thought probes separated by intervals ranging from 40 to 70 s (randomly sampled from a uniform distribution).

### Psychophysiological signal acquisition and preprocessing

#### Electroencephalography and electrocardiography

For the Melbourne dataset, high-density EEG was acquired using 63 EasyCap-mounted active electrodes. A ground electrode was placed over FPz; AFz served as the online reference. The ECG was acquired from electrodes placed on either shoulder; the electro-oculogram (EOG) was recorded from electrodes placed above and below the left and right canthi. All signals were sampled at 500 Hz using a BrainAmp system (Brain Products GmbH) in conjunction with BrainVision Recorder (v1.21.0402; Brain Products GmbH).

For the Paris dataset, EEG records were acquired via 63 actiCap Snap-mounted active electrodes. AFz was again used as the ground electrode, while FCz served as the online reference. The ECG was recorded from an electrode placed below the right shoulder and one placed above the left hip. Horizontal and vertical EOG were acquired from electrodes positioned above and below the dominant eye, and on the left and right outer canthi. All additional electrodes to the EEG (HEOG, VEOG, ECG) were referenced to a ground positioned behind the right shoulder. All signals were sampled at 1000 Hz using BrainVision Recorder (v1.24.0001; Brain Products GmbH).

Offline preprocessing of the raw EEG and ECG was performed in MATLAB R2022a (v9.12.0.1884302) in conjunction with the EEGLAB toolbox (v2024.2.1 ^111^). Data collected in Paris were initially downsampled to 500 Hz to match the online sampling rate in Melbourne. EEG signals were re-referenced to the common average so that the online reference could be reintroduced into the array, and then re-referenced to the average of TP9 and TP10 (in lieu of linked mastoids). The following 62 channels were included in the analysis: Fp1, Fp2, AF7, AF3, AFz, AF4, AF8, F7, F5, F3, F1, Fz, F2, F4, F6, F8, FT9, FT7, FC5, FC3, FC1, FCz, FC2, FC4, FC6, FT8, FT10, T7, C5, C3, C1, Cz, C2, C4, C6, T8, TP7, CP5, CP3, CP1, CPz, CP2, CP4, CP6, TP8, P7, P5, P3, P1, Pz, P2, P4, P6, P8, PO7, PO3, POz, PO4, PO8, O1, Oz, O2.

EEG and ECG records were high-pass filtered (passband edge = 0.5 Hz, transition width = 0.5 Hz) via the ‘pop_eegfiltnew’ function of the ‘firfilt’ plugin (v2.8). Line noise was suppressed at 50 and 100 Hz via the ‘cleanLineNoise’ function from the ‘PrepPipeline’ plugin (v0.55.4 ^112^). The data were then low-pass filtered via ‘pop_eegfiltnew’ (passband edge = 40 Hz, transition width = 10 Hz) and segments between trial blocks excluded. Automated bad channel detection (flatline criterion = 5 s; channel criterion = .8; median number of rejected channels = 0; range = [0, 6]) and artifact subspace reconstruction (burst criterion = 20 ^113^) routines were applied to the EEG via the ‘clean_rawdata’ plugin (v2.91 ^114^). The ECG was z-normalised and R-wave peaks automatically identified using the QRS beat-detection algorithm implemented by the ‘ConvertRawDataToRRIntervals’ function of the PhysioNet Cardiovascular Signal Toolbox (v1.0 ^115^). R-peak annotations were visually inspected (and where necessary, manually corrected) with the aid of the HEPLAB plugin (v1.0.1 ^116^). The ECG signal was then reunited with the EEG and R-peak latency information appended to the event structure.

EEG and ECG data were downsampled to 250 Hz and subjected to independent component analysis (ICA; extended infomax algorithm ^117^). Components were automatically classified using ICLabel (v1.6 ^118^); those classed as ocular, cardiac, or channel noise artefacts with > .90 probability were subtracted from the dataset (median number of rejected components = 4; range = [1, 11]). Data were then subjected to a second round of ICA specifically designed to mitigate the impact of the cardiac field artefact. EEG records were epoched [-200, 200] ms relative to each R-peak and decomposed via ICA. Component and ECG time-series were then low-pass filtered below 25 Hz using a 2-pass, 2nd-order Butterworth filter and Hilbert-transformed to obtain instantaneous phase estimates. Coherence of each component with the ECG signal was calculated as the mean difference between each pair of component-ECG phase angles ^119,120^. Components scoring > 3 S.D. above the subject-level mean coherence were marked as cardiac in nature and subtracted from the full EEG dataset. This procedure was repeated until a maximum of 3 components were excluded, or until no remaining coherence values exceeded the rejection threshold (which was recalculated on each iteration; median number of rejected components = 2; range = [0, 3]). For similar approaches, see Banellis and Cruse ^121^ and Buot and colleagues ^122^.

#### Pupillometry

Eye-movements and pupil size of one eye were sampled at 1000 Hz via an EyeLink 1000 eye-tracking system (SR Research Ltd., Mississauga, Canada). The eye-tracker was calibrated at the beginning of each experimental session using the EyeLink acquisition software. EyeLink 1000 Edf files were imported into MATLAB via the Edf2Mat Toolbox (v1.21.0 ^123^) and preprocessed using custom-built functions based on procedures previously reported by van Kempen and colleagues ^124^ and Corcoran and colleagues ^125^. Blink timings (automatically identified by the EyeLink acquisition software) were corrected via linear interpolation over the blink interval. The start and end points of the interpolated segment were calculated by taking the average pupil size spanning [−200, −100] ms before and [100, 200] ms after the blink period, respectively. The corrected signal was then low-pass filtered below 6 Hz using a 2-pass, 4th-order Butterworth filter. Blinks exceeding 2000 ms were marked as missing data, as were data points spanning 100 ms before and after the blink period.

### Data analysis

#### Signal detection theoretic measures

Responses on Go trials were classed as ‘hits’; failures to respond (errors of omission) were classed as ‘misses’. Responses during NoGo trials (errors of commission) were classed as ‘false alarms’; withheld responses were classed as ‘correct rejections’. Hit rate was calculated as the proportion of Go trials that elicited a response within the 10 s time window preceding the onset of experience sampling (‘preprobe epoch’); false alarm rate was analogously calculated as the proportion of NoGo trials that elicited a response (loglinear correction for extreme values applied following Hautus ^126^). Please note, correct rejections preceded and followed by misses were excluded from analysis on the basis that sustained lack of responding across Go-NoGo-Go trial sequences is indicative of a period of task disengagement (rather than selective response inhibition).

Sensitivity to trial type (Go vs. NoGo) was quantified with the discriminability index *d’*, which was obtained by subtracting z-scored false alarm rates from corresponding z-scored hit rates. A *d′* of 0 indicates chance-level discrimination of Go trials from NoGo trials; more positive values indicate increasing sensitivity. Response bias (decision criterion) was quantified as *c*, the mean of z-scored hit and false alarm rates. A *c* of 0 indicates no bias towards either stimulus category; more positive values indicate greater bias towards issuing a response (i.e., liberal criterion), while more negative values indicate greater bias towards withholding a response (i.e., conservative criterion).

#### Response speed

Response speed estimates were calculated by inverting response times obtained from Go trials (response speed was favoured on account of rendering a more Gaussian-distributed set of observations than response time). Responses registered within 300 ms of trial onset were discarded, since they conflate fast responses to the present trial with (very) slow responses to the preceding trial (see, e.g., Cheyne and colleagues ^127^). Mean response speed (RS*μ*) was obtained by averaging response speed estimates within each preprobe epoch per participant. A minimum of 5 valid responses was required to estimate RS*μ* for a given epoch (median number of rejected epochs = 3; range = [0, 39]). Response speed time-series were then linearly interpolated to impute missing or withheld trial responses, and the coefficient of variation (RS*c_v_*) for each preprobe epoch obtained by dividing the standard deviation of the interpolated series (RS*σ*) by the corresponding mean (RS*μ*):

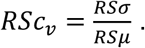

#### Pupil size

Preprocessed pupil size estimates from each set of trials were z-transformed on the subject-level. Estimates spanning the 10 s interval immediately preceding each set of thought probes were averaged to yield preprobe estimates of mean pupil size. Preprobe epochs containing < 5 s of pupil size data were excluded from analysis (median number of rejected epochs = 0; range = [0, 24]).

#### Cardiac parameters

Interbeat intervals (IBIs) – an estimator of cardiac cycle duration (i.e., heart period) – were calculated as the difference between successive R-peaks. IBIs < 300 ms or > 2000 ms in duration were excluded as physiologically implausible. IBIs were z-transformed on the subject-level, and scores exceeding +\-5 z.u. excluded from analysis (on the basis such extreme outliers were likely artifacts due to missing or spurious R-peak annotations; median number of rejected IBIs = 2; range = [0, 30]). The corrected IBI time-series (i.e., tachogram) was then upsampled to 5 Hz and cubic-spline interpolated. Analogous to the reaction speed analysis, mean IBI (IBI*μ*) and coefficient of variation (IBI*c_v_*) estimates were calculated for each preprobe epoch. IBI*c_v_* was natural log-transformed as follows:

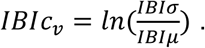

#### Heartbeat-evoked potentials

Heartbeat-evoked potentials (HEPs) were extracted by first applying lowpass filters to the ICA-corrected EEG and ECG signals (passband edge = 20 Hz, transition width = 5 Hz) and segmenting [-0.3, 0.9] s around each R-peak resolved in the 10 s preprobe epoch. R-peaks followed by IBI durations < 600 ms were discarded, and a minimum of 5 R-peaks was required to estimate the HEP for each individual epoch (these criteria resulted in the exclusion of 8 preprobe epochs from one participant and 1 preprobe epoch from another). Epoched data were not ‘baseline-corrected’, since this procedure may contaminate HEP estimates with fluctuations deriving from the preceding cardiac cycle ^1,68^.

#### Mutual information between cardiac and brain signals

Coupling between the EEG and ECG time-series was quantified using mutual information (MI), an information-theoretic measure of the statistical relationship between two random variables^128,129^. MI was estimated using the Gaussian-Copula technique (see Ince and colleagues ^106^) as implemented via v0.4 of the open-source MATLAB code available from https://github.com/robince/gcmi.

ICA-corrected EEG-ECG records were downsampled to 50 Hz and epoched [-16, 2] s relative to the onset of each set of probes. We then used the Nonlinear Mode Decomposition (NMD) toolbox (v2.00 ^130^) to calculate the wavelet transform of the ECG signal between the frequencies [0.5, 2.0] Hz, from which the dominant oscillatory component (i.e., instantaneous heart frequency – a time-resolved representation of HRV) was extracted ^18,131^. EEG signals were bandpass filtered into the following 8 log-spaced low-frequency bands: [0.8, 1.2], [1.2, 1.8], [1.8, 2.6], [2.6, 3.9], [3.9, 5.8], [5.8, 8.6], [8.6, 12.8], and [12.7, 19.0] Hz. Each filtered time-series was Hilbert-transformed to obtain its analytic signal, the real and imaginary components of which were scaled by their corresponding absolute values prior to copula-normalisation (‘copnorm’ function). Normalised time-series were then segmented into 10 s epochs according to the scheme outlined below and the MI between each EEG-ECG channel pair estimated via the ‘mi_gg’ function.

To assess the extent to which coupling dynamics were driven by top-down (brain-to-heart) or bottom-up (heart-to-brain) connectivity, paired signals were offset from one another such that one signal corresponded to the 10 s preprobe epoch while the other preceded this period by some arbitrary time constant. For brain-to-heart coupling, cardiac phase estimates lagged the EEG phase series by the following 12 log-spaced intervals: 159 ms, 204 ms, 262 ms, 337 ms, 433 ms, 557 ms, 715 ms, 920 ms, 1182 ms, 1520 ms, 1954 ms, 2512 ms. For heart-to-brain coupling, EEG phase estimates lagged the cardiac phase series by the following 12 log-spaced intervals: 100 ms, 126 ms, 159 ms, 200 ms, 251 ms, 316 ms, 398 ms, 501 ms, 631 ms, 794 ms, 1000 ms, 1259 ms. Longer latencies were favoured when estimating fluctuations in brain-to-heart coupling due to time delays in the modulation of cardiac pacing incurred at the sino-atrial node ^132^. MI statistics were averaged across time-lags to provide an estimate of brain-to-heart (MI_BRAIN-HEART_) and heart-to-brain (MI_HEART-BRAIN_) coupling for each channel x frequency combination in each preprobe epoch.

#### Statistical analysis

Statistical analysis of behavioural, pupillometric, and cardiac parameters was performed in *R* (v4.2.1 [2022-06-23 ucrt] ^133^) using the *RStudio Desktop* IDE (v2022.07.2+576 ^134^) in conjunction with the ‘tidyverse’ package (v1.3.2 ^135^). These analyses were conducted under a generalised additive mixed-effects model framework (a semiparametric extension of linear mixed-effects modelling) implemented via the ‘mcgv’ package (v1.8-40 ^136,137^). Model diagnostics were assessed using functions from the ‘itsadug’ (v2.4.1 ^138^) and ‘mgcViz’ (v0.1.9 ^139^) packages; model visualisation was accomplished with the aid of ‘cowplot’ (v1.1.1 ^140^), ‘emmeans’ (v1.7.5 ^141^), ‘ggeffects’ (v1.1.2 ^142^), and ‘ggh4x’ (v0.2.8 ^143^).

All models were fitted using restricted maximum likelihood via the ‘gam’ function. Attentional state reports were modelled as unordered categorical data using multinomial logistic regression; vigilance reports were modelled as ordered categorical data. Hit and false alarm rates were modelled using the logit-linked beta distribution; response speed variability was modelled using the log-linked tweedie distribution. All other models assumed Gaussian-distributed errors. Dependent variables were regressed onto a covariate encoding stimulus type (Digit vs. Face) and the independent variable of interest (except in the case of IBI*μ* and IBI*c_v_*, which were included together as orthogonal predictors). When models were fitted to data from Melbourne and Paris, site of data collection was included as an additional covariate. Models were additionally equipped with a by-participant factor smooth on probe number to control for time-on-task and reduce the temporal autocorrelation of residuals (see Baayen and colleagues ^144^).

Brain-heart connectivity measures were analysed in MATLAB R2022a (v9.12.0.1884302) using a mass linear mixed-effects modelling approach ^145^. Linear mixed-effects models including fixed effects of attentional state or vigilance level, covariates for stimulus type and site, and random intercepts for participant identity, were fitted to HEP or MI estimates using maximum likelihood estimation (‘fitlme’). HEP models additionally included covariates for IBI*μ* and IBI*c_v_*. Marginal estimates for each channel x time-point (HEP) or channel x frequency-band (MI) combination were extracted for subsequent mass univariate linear regression analysis (‘lmeEEG_regress’). T-values obtained from these regression models were then compared against those generated from 1000 random permutations of the design matrix (‘lmeEEG_permutations’) using threshold-free cluster enhancement ^146,147^ (TFCE; implemented via ‘lmeEEG_TFCE’). For the HEP analysis, the time-window of interest ranged [200, 600] ms post R-peak.

As a further precaution against the possibility of spurious results marring our HEP analysis, t-values from significant clusters resolved using TFCE were compared to those generated by replicating the mass univariate analysis on surrogate R-peaks ^73,148^. We created surrogate R-peaks by randomly permuting each IBI time-series within the 40 s period preceding each set of thought probes and time-locking our HEP analysis to the latencies implied by the reordered IBIs. This procedure was repeated 100 times to build a null distribution of surrogate t-values that reflected differences in EEG fluctuations that were not systematically related to the heartbeat (since the temporal dependence between EEG responses and cardiac systole had been broken) ^149^. Observed t-values that failed to exceed the permutation threshold defining a Monte Carlo p-value < .05 (two-tailed) were masked as non-significant.

### Data and code availability

The datasets analysed in this report can be accessed from the Open Science Framework platform (Melbourne: https://osf.io/ey3ca/ ^150^; Paris: https://osf.io/v9xsw/ ^151^). The code used to perform these analyses is available on GitHub: https://github.com/corcorana/WIM_HB ^152^.

## Results

### Prevalence of disengaged states and impact on behavioural performance

Similar to the original analysis reported by Andrillon and colleagues ^107^, participants tested in Melbourne declared being on-task an average 49% of times probed; mind-wandering and mind-blanking were reported an average 39% and 13% of the time, respectively. A comparable profile of responses was observed in the Paris dataset (ON: 56% probes; MW: 30% probes; MB: 14% probes). These findings are broadly consistent with previous experience-sampling research examining on-task vs. disengaged states ^153^. One participant in the Melbourne dataset and three in the Paris dataset reported being on-task throughout the session; hence, these individuals were excluded from analyses involving attentional state reports (but were retained otherwise).

Mind-wandering and mind-blanking states were reported more frequently with time-on-task in both datasets (MW_MEL_: β = 0.46, s.e. = 0.06, z = 7.16, p < .001; MW_PAR_: β = 0.32, s.e. = 0.05, z = 6.33, p < .001; MB_MEL_: β = 0.35, s.e. = 0.09, z = 3.79, p < .001; MB_PAR_: β = 0.15, s.e. = 0.07, z = 2.27, p = .024). They were also associated with lower vigilance ratings (MW_MEL_: β = −0.87, s.e. = 0.11, z = 7.64, p < .001; MW_PAR_: β = −0.83, s.e. = 0.08, z = 10.95, p < .001; MB_MEL_: β = −1.28, s.e. = 0.17, z = 7.66, p < .001; MB_PAR_: β = −1.36, s.e. = 0.11, z = 12.28, p < .001) and smaller pupil size (MW_MEL_: β= −0.34, s.e. = 0.11, z = 3.24,, p = .001; MW_PAR_: β= −0.25, s.e. = 0.08, z = 3.16., p = .002; MB_MEL_: β = −0.54, s.e. = 0.14, z = 3.79, p < .001; MB_PAR_: β = −0.50, s.e. = 0.11, z = 4.61, p < .001), than on-task reports.

Using signal detection theory ^154,155^, we calculated measures of sensitivity (*d’*) and response bias (*c*) in 10 s trial epochs preceding thought probes. Sensitivity was significantly decreased in epochs preceding off-task reports (MW_MEL_: β = −0.26, s.e. = 0.04, t = 5.92, p < .001; MW_PAR_: β= −0.34, s.e. = 0.04, z = 8.94., p < .001; MB_MEL_: β = −0.32, s.e. = 0.06, t = 5.13, p < .001; MB_PAR_: β = −0.46, s.e. = 0.05, z = 9.27, p < .001) as compared to on-task reports. Bias was significantly increased (i.e., increased tendency to produce false alarms) in epochs preceding mind-wandering (MW_MEL_: β = 0.09, s.e. = 0.02, t = 4.21, p < .001; MW_PAR_: β = 0.06, s.e. = 0.02, t = 3.29, p = .001), but not mind-blanking (MB_MEL_: β = 0.02, s.e. = 0.03, t = 0.65, p = .516; MB_PAR_: β = 0, s.e. = 0.02, t = 0.02, p = .982).

Mean response speed (1/Response Time; RS*μ*) on Go trials preceding mind-wandering reports did not significantly differ from on-task reports (MW_MEL_: β = 0, s.e. = 0.01, t = 0.03, p = .977; MW_PAR_: β = −0.02, s.e. = 0.02, t = 1.03, p = .304). However, mind-blanking was associated with significant response slowing (MB_MEL_: β = −0.05, s.e. = 0.02, t = 2.63, p = .009; MB_PAR_: β = −0.04, s.e. = 0.02, t = 2.00, p = .046). Response speed variability (RS*c_v_*) was significantly increased prior to both mind-wandering (MW_MEL_: β = 0.06, s.e. = 0.02, t = 2.25, p = .025; MW_PAR_: β = 0.05, s.e. = 0.02, t = 2.03, p = .021) and mind-blanking (MB_MEL_: β = 0.11, s.e. = 0.04, t = 3.02, p = .003; MB_PAR_: β = 0.10, s.e. = 0.03, t = 3.35, p < .001) relative to on-task epochs. See Table 1 for summary statistics.

**Table 1.**
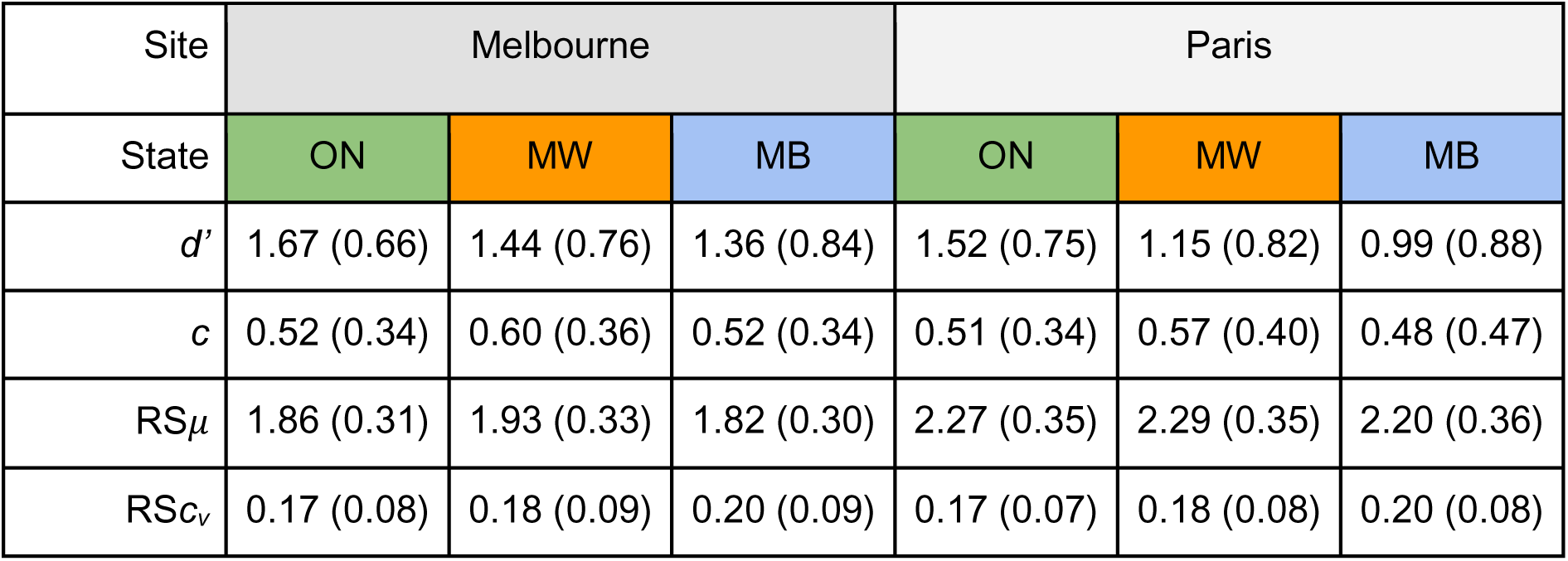
Summary statistics for behavioural measures calculated under each attentional state for both datasets. Mean (standard deviation) for sensitivity (d’), bias (c), response speed (RS*μ*) and coefficient of variation (RSc_v_) measures calculated within each attentional state response category for both the Melbourne and Paris datasets. ON = on-task; MW = mind-wandering; MB = mind-blanking.

| Site | Melbourne |  |  | Paris |  |  |
| --- | --- | --- | --- | --- | --- | --- |
| State | ON | MW | MB | ON | MW | MB |
| $d'$ | 1.67 (0.66) | 1.44 (0.76) | 1.36 (0.84) | 1.52 (0.75) | 1.15 (0.82) | 0.99 (0.88) |
| $c$ | 0.52 (0.34) | 0.60 (0.36) | 0.52 (0.34) | 0.51 (0.34) | 0.57 (0.40) | 0.48 (0.47) |
| $RS_{\mu}$ | 1.86 (0.31) | 1.93 (0.33) | 1.82 (0.30) | 2.27 (0.35) | 2.29 (0.35) | 2.20 (0.36) |
| $RSc_v$ | 0.17 (0.08) | 0.18 (0.09) | 0.20 (0.09) | 0.17 (0.07) | 0.18 (0.08) | 0.20 (0.08) |

Overall, these results confirm previous reports from overlapping and different datasets ^87,88,101,156^ showing distinct signatures of mind-wandering and mind-blanking on cognitive performance. Given the remarkable consistency of the findings observed across the Melbourne and Paris datasets, the remainder of this report focuses on a single set of results obtained from analysis of the combined dataset (please note, site of data collection was included as a covariate within each model).

### Cardiac measures afford complementary information about arousal, attentional, and behavioural states

Having confirmed the expected differences in behavioural and pupillometric measures across attentional and vigilance state reports, we next sought to examine whether cardiac parameters derived from epochs preceding thought probes predict attentional and arousal states and their behavioural correlates.

Mixed-effects modelling revealed a positive association between mean interbeat interval (IBI*μ*) and off-task attentional states: increased IBI*μ* (i.e., lower heart-rate) significantly increased the propensity to report both mind-wandering (β = 0.31, s.e. = 0.07, z = 4.22, p < .001) and mind-blanking (β = 0.70, s.e. = 0.10, z = 7.08, p < .001). Moreover, increased IBI*μ* was associated with lower alertness ratings (β = −0.35, s.e. = 0.07, z = 4.89, p < .001) and smaller pupil size (β = −0.21, s.e. = 0.02, z = 11.42, p < .001), consistent with a global shift towards parasympathetic dominance and hypoarousal ^157,158^. HRV (interbeat interval coefficient of variation; IBI*c_v_*) did not significantly predict mind-blanking (β = 0.12, s.e. = 0.08, z = 1.57, p = .117) nor vigilance level (β = −0.05, s.e. = 0.05, z = 0.95, p = .035), but did predict increases in mind-wandering (β = 0.10, s.e. = 0.05, z = 1.99, p = .046) and pupil dilation (β = 0.07, s.e. = 0.01, z = 5.64, p < .001; see Figure 2).

**Figure 2:**
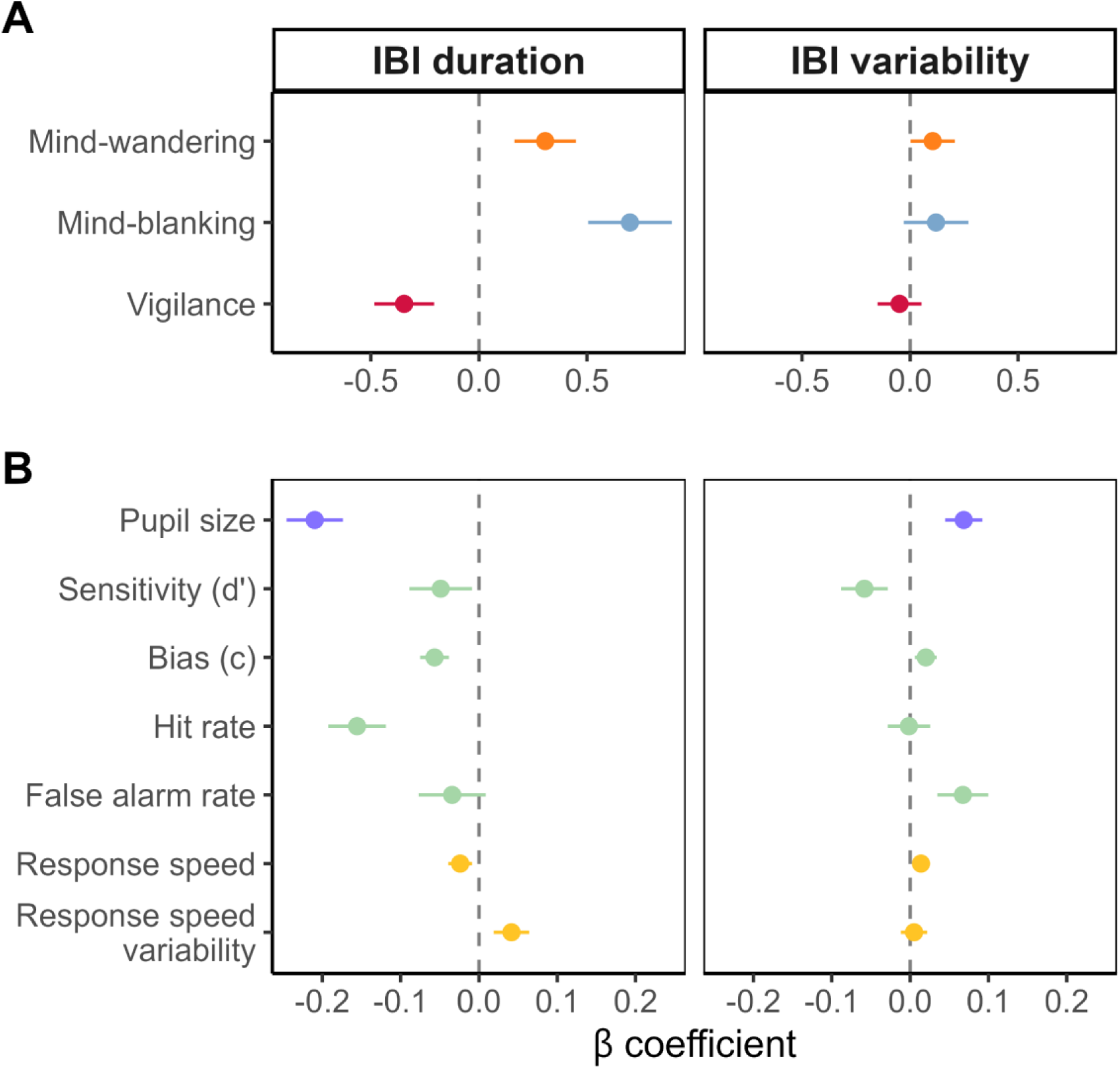
**Summary of modelled relationships between cardiac parameters and attentional, arousal, and behavioural state variables** A: Estimated effect of a z-unit increase in mean interbeat interval (IBI) duration (left) and coefficient of variation (right) on subjective reports of attentional state and vigilance level. Mind-wandering and mind-blanking coefficients encode differences relative to the on-task state, where a positive coefficient indicates an increase in the likeliness of reporting mind-wandering/mind-blanking (per z-unit increase in IBI duration/variability). B: Estimated effect of a z-unit increase in mean IBI duration (left) and coefficient of variation (right) on mean pupil size, signal detection theoretic and response speed measures. Error bars represent 95% confidence intervals. All estimates derived from 10 s epochs immediately preceding thought probes.

Turning next to behavioural performance during the SART, a negative association was observed between IBI*μ* and *d’* (β = −0.05, s.e. = 0.02, t = 2.40, p = .027). This effect was driven by a reduction in hit rate as IBI*μ* increased (β = −0.16, s.e. = 0.02, t = 8.30, p < .001). Increased IBI*μ* was also associated with decreased *c* (β = −0.06, s.e. = 0.01, z = 6.03, p < .001), RS*μ* (β = −0.02, s.e. = 0.01, z = 3.11, p = .002), and increased RS*c_v_* (β = 0.04, s.e. = 0.01, z = 3.59, p < .001), but did not predict false alarm rate (β = −0.03, s.e. = 0.02, z = 1.57, p = .117). IBI*c_v_* was likewise negatively associated with *d’* (β = −0.06, s.e. = 0.02, t = 3.84, p < .001); however, this effect was driven by an increase in false alarm rate (β = 0.07, s.e. = 0.02, z = 4.05, p < .001). IBI*c_v_* also predicted increases in *c* (β = 0.02, s.e. = 0.01, t = 2.82, p = .005) and RS*μ* (β = 0.01, s.e. = 0.01, t = 2.46, p = .014), but not hit rate (β = 0, s.e. = 0.01, z = 0.11, p = .910) or RS*c_v_* (β = 0.01, s.e. = 0.01, z = 0.58, p = .559).

### Error-related cardiac deceleration does not account for differences across attentional states

Since mind-wandering and mind-blanking reports were associated with decrements in SART performance, we considered whether the differences we observed in IBI*μ* between task-focused and disengaged states could be attributed to increased rates of erroneous responding during periods of task disengagement. IBIs are known to lengthen in response to performance errors even in the absence of explicit feedback ^46^; hence, increased false alarm rates might be expected to elicit more frequent bouts of cardiac deceleration, thereby lowering IBI*μ*.

To assess this possibility, we identified all NoGo trials occurring within each 10 s preprobe epoch and extracted the z-scored duration of the 3 IBIs preceding and following the interval in which the NoGo stimulus was presented (IBIs following the NoGo stimulus were discarded if they were interrupted by thought probes). IBI duration was then regressed onto factors encoding the ordinal position of each IBI relative to NoGo trial onset, behavioural response category (CR vs. FA), and subsequent attentional state report (ON vs. MW vs. MB), along with interactions between these factors. Covariates for stimulus type (Digit vs. Face), site of data collection (Melbourne vs. Paris), and random factor smooths for probe and participant identity were also included, consistent with the models reported in the previous two sections.

Model predictions are visualised in Figure 3. As expected, the main effect of IBI order was significant (F(6) = 27.82, p < .001); the first IBI following NoGo trial onset was significantly longer than both the preceding interval (IBI_+1_-IBI_0_ = 0.15, se = 0.02, t = 7.57, p_adj_ < .001) and the subsequent interval (IBI_+1_-IBI_+2_ = 0.12, s.e. = 0.02, t = 5.75, p_adj_ < .001), consistent with the occurrence of a cardiac deceleration response to salient (i.e., rare or task-relevant) stimuli. The main effect of attentional state was also significant (F(2) = 51.71, p < .001), consistent with the observation that IBIs tend to be longer on average in epochs preceding mind-wandering (β = 0.06, s.e. = 0.02, t = 3.85, p < .001) and mind-blanking reports (β = 0.22, s.e. = 0.02, t = 10.13, p < .001) than on-task reports. Interestingly, IBIs tended to be longer in sequences spanning a correct rejection (F(1) = 23.95, p < .001), irrespective of the subsequent attentional state report. This observation is consistent with our finding that increased IBI*μ* predicted decreased response bias. Moreover, false alarms did not generate more profound cardiac decelerations than those observed when responses to NoGo trials were withheld. Together, these observations indicate that the increase in IBI*μ* associated with mind-wandering and mind-blanking was unlikely to be driven by higher error rates of erroneous responding.

**Figure 3:**
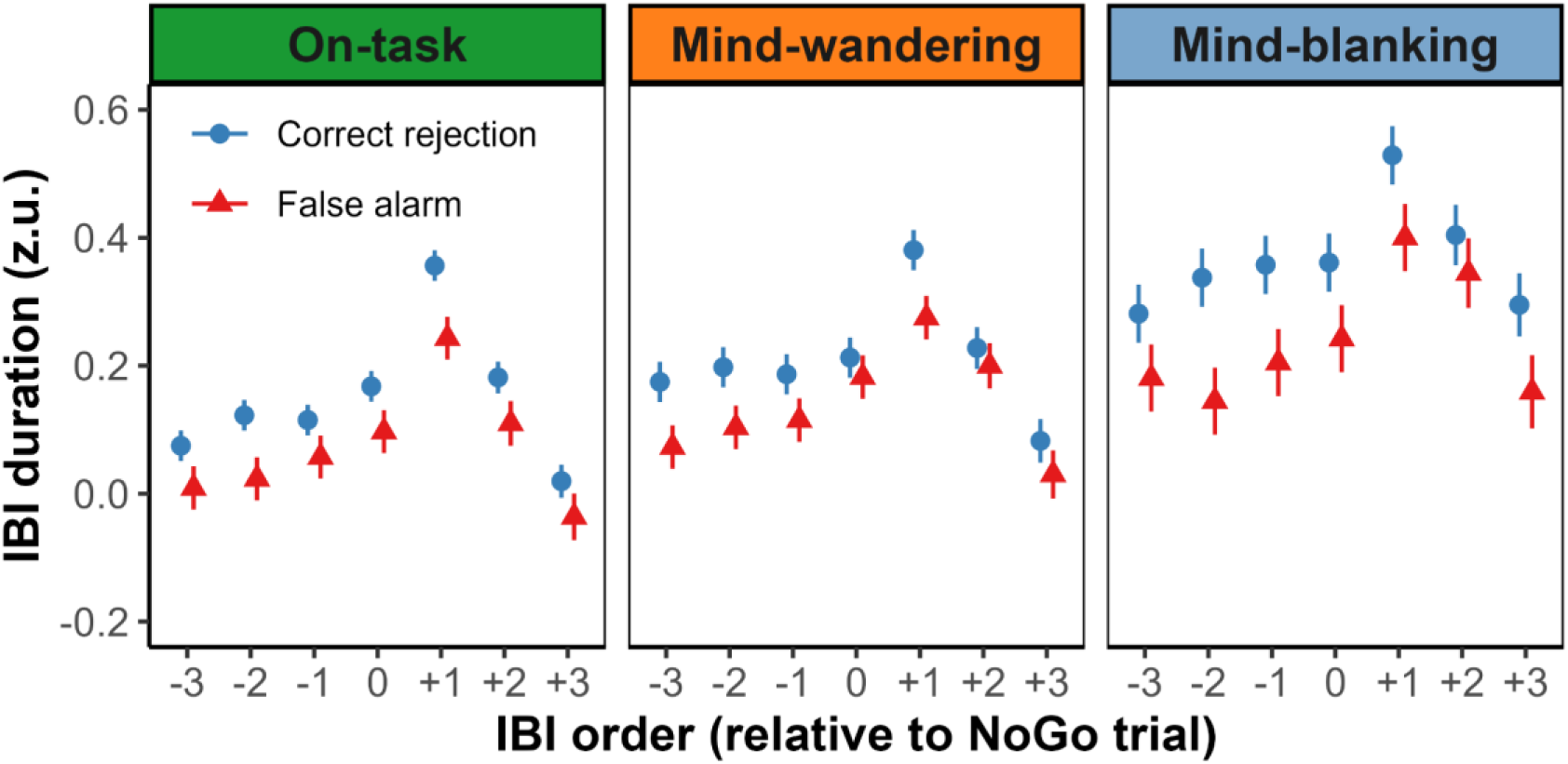
**Interbeat intervals show similar NoGo response profiles across attentional states.** Estimated marginal mean IBI duration (z-units) for IBI sequences centred on IBIs containing the onset of NoGo trials within the 10 s preprobe epoch. IBI sequences featuring NoGo trials that elicited correct rejections are denoted as blue circles; those associated with false alarms (errors of commission) are denoted as red triangles. Estimates are further faceted according to the attentional state that was reported following the epoch. Error bars represent the standard error of the mean (SEM).

### Heartbeat-evoked potentials are modulated by mind-wandering

To assess whether task-disengagement is accompanied by altered interoceptive processing of cardiac signals, we investigated how neural responses time-locked to the heartbeat (i.e., heartbeat-evoked potentials; HEPs) varied as a function of attentional state and vigilance level reports. HEPs were estimated on the participant-level for R-peaks captured within the 10 s epoch preceding each set of thought probes. ERPs were also computed for the corresponding ECG signal. Linear mixed-effects models were fitted for each channel x time-point combination in order to examine the impact of attentional states and vigilance ratings on the time course of the ECG-ERP (Figure 4A) and HEP (Figure 4B). To correct for multiple comparisons, we applied a cluster permutation approach across all channels for the time-window of interest (200–600 ms post R-peak). As an additional safeguard against spurious results, t-values from significant clusters were checked against those derived from a set of surrogate R-peak latencies (see Methods for further details).

**Figure 4:**
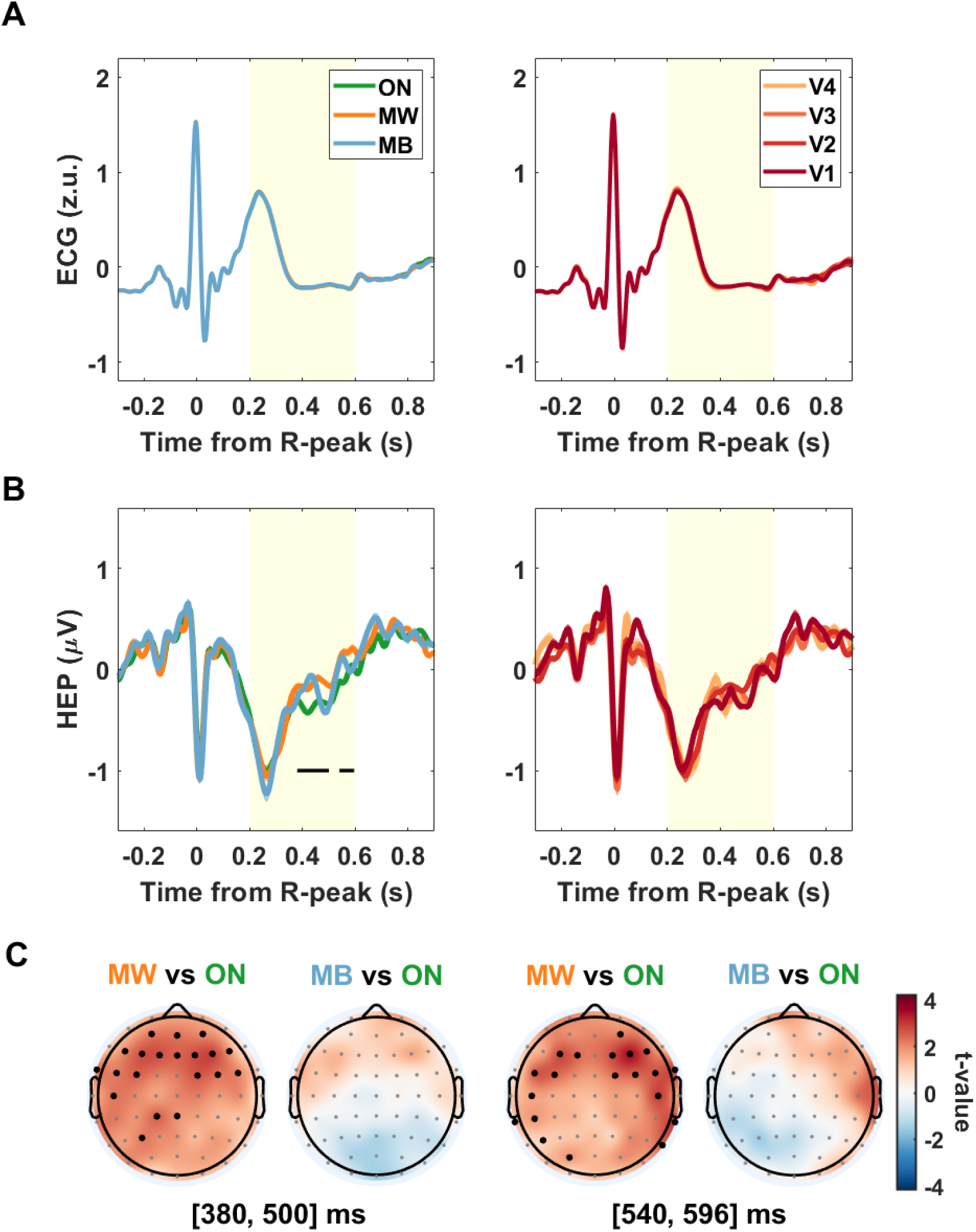
**Heartbeat-evoked potentials are modulated by attentional state but not vigilance level.** A: Event-related potential (ERP) of the electrocardiogram (ECG) averaged across individuals for the different attentional states (left; ON = on-task, MW = mind-wandering, MB = mind-blanking) and vigilance levels (right; V1 = extreme sleepiness, V4 = extreme alertness). B: Heartbeat-evoked potential (HEP) for the different attentional states (left) and vigilance levels (right). Each HEP was averaged across the 11 channels that revealed significant differences between mind-wandering and on-task states (MW vs. ON) in both the earlier and later clusters. For both A and B, shaded areas denote the standard error of the mean (SEM); yellow shading indicates time-window of interest; horizontal black bars indicate the timespan of significant clusters in the MW vs. ON contrast (p_cluster_< 0.05, corrected for surrogate t-values; see Methods). C: Topographic representation of t-values from the attentional state contrasts averaged over the timespan of the two significant clusters marked in B. Black dots indicate electrodes that exceeded both the permutation-test and surrogate t-value thresholds for cluster membership (see Methods).

Following these procedures, we found two significant clusters indicative of modulations of the HEP during epochs associated with mind-wandering (relative to those observed prior to on-task reports). The first cluster spanned [380, 500] ms, and included frontal and left-parietal scalp regions; the second cluster spanned [540, 596] ms, and evinced a more lateralised topography (Figure 4C). These fluctuations were not accompanied by corresponding differences in the time-course of ECG-ERP (Figure 4A), suggesting that late modulation of the HEP during mind-wandering states derived from genuine differences in cortical responses to the heartbeat. No significant clusters were identified when HEPs preceding mind-blanking reports were contrasted with those preceding on-task reports, nor when HEPs were compared across different vigilance levels (see Figure 4B for representative ERPs).

### Brain-heart dynamics decouple with task-disengagement and fatigue

Finally, we sought to complement our analysis of cortical responses to the heartbeat in the time-domain by investigating how the evolution of cardiac and brain activity in the frequency-domain varies across attentional states and vigilance levels. To this end, we estimated the mutual information (MI) shared between the instantaneous phases of the dominant heart frequency and low-frequency EEG signals within each preprobe epoch. Computing this measure at various time-lags enabled us evaluate how MI changed depending on which time-series led the other, whereby changes in coupling strength were assumed to be driven by the earlier-occurring signal (e.g., when the EEG preceded the ECG signal [BRAIN-HEART], increased MI was interpreted as evidence of increased top-down cardiac regulation; when the ECG preceded the EEG signal [HEART-BRAIN], increased MI was interpreted as evidence of increased interoceptive processing of bottom-up input). MI estimates were averaged across time-lags within each set of directionalities, and subjected to a similar mixed-effects modelling and cluster permutation procedure to that employed in our HEP analysis (see Methods for further details).

Very similar profiles of time-averaged MI were resolved across frequencies in both the brain-to-heart and heart-to-brain directionality (Figure 5A). MI tended to be highest for the lowest two EEG frequency bands included in the analysis (0.8–1.8 Hz), which correspond to the lower half of the canonical delta band (1–4 Hz). MI declined by approximately an order of magnitude in the transition from (high) delta (1.8–3.9 Hz) to alpha band activity (8.5–12.8 Hz), and remained low when estimated for the (low) beta band (12.7–19.0 Hz).

**Figure 5:**
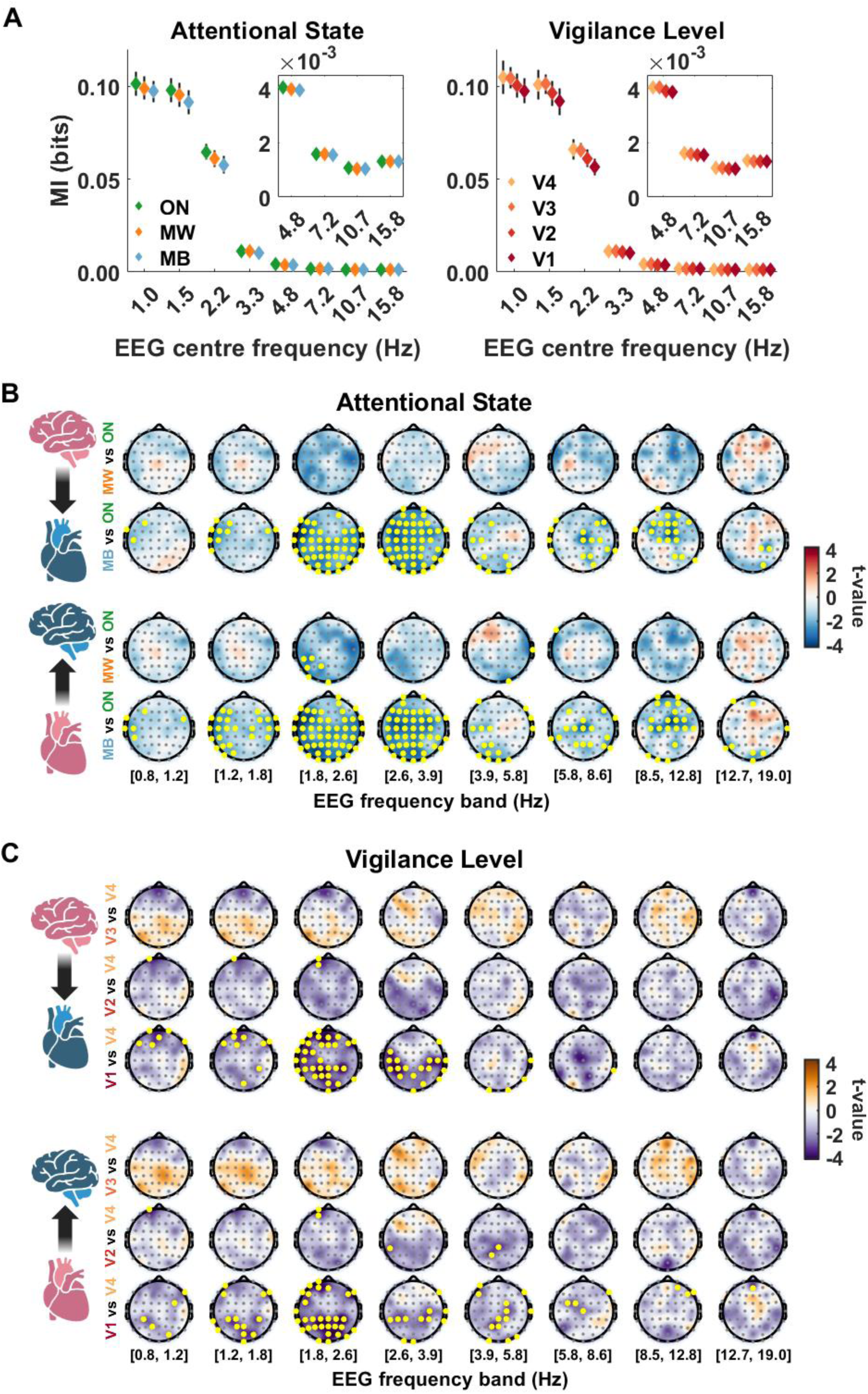
**Phase coupling between brain and heart signals decreases with task-disengagement and sleepiness.** A: Mutual information (MI) profiles averaged across channels and directionalities for attentional state (left; ON = on-task, MW = mind-wandering, MB = mind-blanking) and vigilance level (right; V1 = extreme sleepiness, V4 = extreme alertness). Error bars indicate the standard deviation of channel-wise MI estimates. Inset panels depict averaged MI estimates for higher frequency EEG bands at increased magnification. B: Topographic representation of t-values from the attentional state contrasts for each frequency band and directionality (brain-to-heart top; heart-to-brain bottom). Yellow dots indicate electrodes that exceeded the permutation test t-value threshold. C: Topographic representation of t-values from the vigilance level contrasts for each frequency band and directionality (brain-to-heart top; heart-to-brain bottom). Yellow dots indicate electrodes that exceeded the permutation test t-value threshold.

Focusing on differences in MI_BRAIN-HEART_ across attentional states (Figure 5B), our analysis revealed no significant changes in brain-to-heart phase coupling between on-task and mind-wandering epochs. By contrast, mind-blanking epochs revealed significant decreases in MI_BRAIN-HEART_ estimates relative to on-task epochs. This effect was evident to varying degrees across all included frequency bins, but most spatially extensive in the bands spanning [1.8, 3.9] Hz. Significant clusters were also resolved at higher frequencies, including a left-temporal and right fronto-parietal cluster in the [5.8, 8.6] Hz band, and a predominantly frontal cluster in the [8.5, 12.8] Hz band. Repeating this analysis on MI_HEART-BRAIN_ contrasts revealed a qualitatively similar pattern of results, albeit with a few additional significant effects. A significant left-parietal cluster was resolved for mind-wandering epochs in the [1.8, 2.6] Hz band. Mind-blanking epochs evinced more extensive left- and right-sided clusters in the [1.2, 1.8] Hz band as compared to the MI_BRAIN-HEART_ contrast for the same band. Large clusters were again resolved in the [1.8, 3.9] Hz range, while broadly similar effects were observed in the higher frequency bands.

Turning next to vigilance reports, we compared how brain-heart phase coupling fluctuated relative to states of ‘extreme alertness’ (V4; Figure 5C). We found no significant effects for either directionality when contrasting MI estimates from epochs preceding V3 reports to those preceding V4; however, lower levels of alertness were associated with significant decreases in brain-to-heart phase coupling. Moderate sleepiness (V2) was associated with significant decreases in MI_BRAIN-HEART_ and MI_HEART-BRAIN_ across 2 frontal channels in the [1.8, 2.6] Hz band; MI_HEART-BRAIN_ was also significantly reduced across a small number of left-parietal channels in the [2.6, 5.8 Hz] range. Extreme sleepiness (V1) was accompanied by more spatially extensive reductions in brain-heart phase coupling, where the largest clusters spanned the [1.2, 3.9] Hz range. Clusters tended to include more frontal electrode channels in the brain-to-heart analysis than the corresponding heart-to-brain contrasts. Unlike the mind-blanking effects, few significant effects were observed for either directionality in frequencies above 5.8 Hz.

## Discussion

This study investigated how spontaneous fluctuations of attention are related to subjective and physiological measures of arousal, with specific focus on the brain-heart axis ^60^. In summary, we found that task disengagement (i.e., self-reported episodes of mind-wandering or mind-blanking) was characterised by decreases in autonomic arousal, as indicated by reductions in prevailing heart-rate (increased IBI*μ*) and smaller pupil size. These physiological effects manifested subjectively as increased sleepiness and behaviourally as reduced sensitivity (*d’*) and increased response speed variability (RS*c_v_*). Notably, mind-blanking states were accompanied by more pronounced reductions in heart-rate and response speed (RS*μ*) than mind-wandering states, thus suggesting that mind-blanking is more likely to occur in the context of hypoarousal. Mind-wandering, by contrast, was characterised by subtler decreases in arousal that were associated with more stereotypic behaviour (increased response bias (*c*), unchanged response speed) and increased heart-rate variability (HRV; as quantified by IBI*c_v_*). Independent of attentional state, greater HRV predicted increased pupil dilation, response bias, response speed, and false alarm rate, suggesting transient fluctuations in heartbeat dynamics afford unique information pertaining to what appear to be impulsive patterns of behaviour.

Having identified how cardiac parameters vary in relation to subjective and objective indices of task focus and disengagement, we next sought to examine whether such fluctuations are accompanied by distinctive profiles of brain-heart interaction. We turned first to the heartbeat-evoked potential (HEP), an extensively-studied psychophysiological marker of interoceptive processing that has been associated with the focus of attention ^67^. This analysis revealed a late modulation of HEPs obtained in the period preceding mind-wandering reports, relative to those obtained prior to on-task reports. No differences were observed between mind-blanking and on-task states, or amongst vigilance levels. Finally, we used a mutual information (MI) based method to estimate phase coupling between brain and heart signal components. This analysis revealed evidence of decreasing MI as participants transitioned from task-focused to disengaged states (especially mind-blanking), and as vigilance level declined. Together, these findings suggest that more profound states of hypoarousal are accompanied by a generalised decoupling of brain and heart dynamics, paralleling the decoupling from exteroceptive inputs reported elsewhere ^159^.

Our results add to recent evidence that mind-blanking constitutes a low-arousal state distinct from other forms of task disengagement, such as mind-wandering ^87,88,159–161^. Our focus on cardiac parameters and measures of brain-heart coupling complement previous reports on the oscillatory, event-related, and information-theoretic EEG signatures of mind-blanking ^87,159^, demonstrating that these signatures are accompanied by detectable alterations in peripheral electrophysiology and cerebro-peripheral connectivity. They also cohere with recent evidence that mind-blanking reports elicited during task-free resting-state are more prevalent in the context of low-arousal conditions induced by sleep deprivation ^88^. While the latter study indicated that the addition of cardiac parameters to EEG measures helped improve the automated classification of mental state reports, our findings isolate the information afforded by particular cardiac parameters with respect to subjective, neurophysiological, and behavioural measures of arousal and attentional state dynamics.

Mean interbeat interval duration (IBI*μ*) and pupil size provide complementary psychophysiological indices of arousal state that reflect the balance between sympathetic and parasympathetic tone ^157,162–164^. Our findings reveal that mind-wandering and mind-blanking episodes are characterised by a progressive shift towards parasympathetic dominance, as indicated by corresponding decreases in prevailing heart-rate and pupil size. This finding is consonant with the reduction in heart-rate observed during the transition from wakefulness to deep sleep ^165–168^ – a phenomenon tied to arousal state modulation (not just circadian variability) ^169,170^. Notably, the effects we report here were observed after statistically controlling for time-on-task (i.e., by specifying factor-smooths on probe number for each participant), indicating that markers of parasympathetic outflow afford specific information (related to cardiac activity) about the transition from on-task through to mind-blanking states over and above generic effects pertaining to prolonged task performance (e.g., motivational state, mental fatigue, drowsiness, etc.) ^171–175^.

Intriguingly, short-term heart-rate variability – which is commonly used to measure phasic fluctuations in parasympathetic activity, and which also typically increases in sleep ^176–179^ – was positively associated with mean pupil size. While this finding might seem counterintuitive given the aforementioned negative association between IBI*μ* and pupil size, two points need to be borne in mind: First, increases in HRV need not derive from tonic increases in parasympathetic outflow (or a relative shift towards parasympathetic dominance), but may simply reflect altered phasic patterning of parasympathetic discharge ^162^. Second, by taking the coefficient of variation as our estimator of HRV (and by controlling for potential confounding from time-on-task effects), we ensured that our measures of heartbeat activity were orthogonal to one another – thus enabling us to uncover different relations between unique components of cardiac control.

Increased pupil-linked arousal in the context of enhanced HRV may be mechanistically explained by phasic input from ascending parasympathetic pathways into the noradrenergic locus coeruleus ^180–183^, a major neuromodulatory system responsible for the regulation of arousal and task (dis)engagement ^184,185^. Given that IBI*c_v_* was also associated with increases in response bias, speed, and false alarm rate (errors of commission), the positive relation we observed between pupil dilation and HRV may be partially explained by error-related orienting ^45,186–189^, whereby the unexpected occurrence of an erroneous behavioural action (i.e., responding to a NoGo stimulus) precipitated a transient increase in arousal and the reallocation of attentional resources toward the task at hand ^190–194^. In turn, the positive association between IBI*c_v_* and mind-wandering (but not mind-blanking) suggests that mind-wandering is associated with episodes of low but unstable arousal. Accordingly, mind-wandering could represent transitions to or from low levels of arousal, whereas mind-blanking would represent stable periods of hypoarousal.

Our analysis of neural responses time-locked to the heartbeat revealed a late modulation of HEPs derived from epochs preceding mind-wandering reports. Since these epochs were also characterised by increases in IBI*c_v_*, one potential explanation for this finding is that neural responses to afferent feedback generated by the heartbeat were amplified as a consequence of greater uncertainty about the precise timing of successive bursts of interoceptive input. Notice, however, that we included IBI*c_v_* as a covariate in our regression-based HEP analysis; hence, we believe this effect to be specific to the context of mind-wandering per se, rather than a generic consequence of the temporal uncertainty induced by more variable (i.e., unpredictable) fluctuations in cardio-afferent feedback. Functional brain imaging has revealed the default mode network as an important neural substrate for both mind-wandering ^195–197^ and parasympathetic activity ^61,198^, while self-related cognitive processes typically engaged during episodes of mind-wandering have been linked with altered heartbeat-evoked responses ^73–75^. Collectively, these findings suggest that increases in HRV and HEP amplitude reflect changes in central autonomic activity that stem from the enhanced functional integration of brain regions involved in the spontaneous generation of task-unrelated thought.

In contrast to the circumscribed findings of our HEP analysis, our MI analysis revealed that phase coupling between evolving heart dynamics and bandpass-filtered EEG time-series progressively decreased as participants transitioned from higher to lower states of arousal. This effect is congruent with evidence of cardiac and low-frequency EEG decoupling during the transition from wakefulness to sleep ^199,200^. We observed more widespread decreases in coupling strength in both sensor and frequency space when participants reported mind-blanking (rather than mind-wandering) and – to a lesser extent – extreme sleepiness (as opposed to mild or moderate decreases in alertness). The qualitative similarity evinced by top-down and bottom-up estimates of brain-heart connectivity suggest a functional decoupling of the dynamics expressed by these two systems, rather than a relative shift towards a more predominantly top-down or bottom-up pattern of information exchange. That is, low arousal states appear to be characterised by more intrinsically-driven patterns of cardiac activity, which in turn exert less influence on the evolving dynamics of low-frequency brain activity.

Previous work has suggested that endogenous fluctuations in attentional focus may reflect transient oscillations between external (exteroceptive) vs. internal (interoceptive) states ^19,69,72,201,202^. One might therefore anticipate that task disengagement is associated with stronger brain-heart interactions, on the basis that individuals disengaged from their external environment are consequently more engaged with their internal environment. While this account may hold for certain scenarios, our analysis furnishes evidence against a simple switch-like mechanism whereby reduced attention to the external environment necessarily entails increased interoceptive monitoring and physiological regulation. Indeed, it seems plausible to us that effective task engagement should call for a concerted increase in brain-heart integration in addition to the allocation of attentional resources towards task-relevant stimuli, such that evolving physiological dynamics may be deftly calibrated in accordance with evolving situational demands (e.g., transiently increasing heart-rate to meet the metabolic costs of completing a particularly challenging component of a task). During more quiescent states, by contrast, the brain might take the opportunity to disengage from both its external and internal environments, thereby ‘relaxing’ control over the cardiovascular system (or delegating control to lower-order networks; e.g., those situated within the brainstem and hypothalamus).

In conclusion, this study adds to the growing body of literature distinguishing mind-wandering and mind-blanking as distinctive attentional states. Our analysis of cardiac dynamics in conjunction with subjective, behavioural, and pupillometric measures provides complementary evidence that mind-wandering and mind-blanking arise as arousal levels progressively decrease, likely mediated by global increases in parasympathetic tone. Late modulation of heartbeat-evoked responses further indicates that mind-wandering is associated with amplified cortical processing of interoceptive signals. The allocation of attentional resources towards interoceptive processes that typically unfold outside of awareness may contribute to the subjective experience of losing track of one’s surroundings. Furthermore, decreasing arousal is associated with decoupling of brain and heart dynamics, suggesting task disengagement is accompanied by a concomitant disengagement from the monitoring and regulation of internal bodily states. Finally, the link between heartbeat fluctuations, pupil dilation, and impulsive-like behaviour suggests transient bursts of vagal outflow promote phasic increases of arousal, possibly on account of driving input to the noradrenergic locus coeruleus. Together, these findings highlight how the neural processing and regulation of cardiac activity varies as a function of attentional state, shedding light on the way brain-body interactions shape cognitive and behavioural dynamics subtending multiple timescales.

## Acknowledgements

Funded by the European Union (ERC, SleepingAwake, 101116748). The research leading to these results has received funding from the national program “Investissements d’avenir” ANR-10-IAIHU-0006. Part of this work was carried out in the CENIR of ICM. AWC and JH acknowledge the support of the Three Springs Foundation. TA was supported by a National Health Medical Research Council Ideas Project (APP1183280) and a Long-Term Fellowship from the Human Frontier Science Programme (LT000362/2018-L). ALC is supported by a scholarship from the Fondation pour la Recherche Médicale (FDT202404018709). We thank Teigane Mackay and Devangna Tangri for data collection in Melbourne, Marie Degrave, Laurent Hugueville, and Amandine Carrie for data collection at the Paris Brain Institute, Naotsugu Tsuchiya and Jennifer Windt for their support to TA at the inception of this line of research, and three anonymous reviewers for their feedback on a previous version of this manuscript.

## Conflict of interest statement

The authors declare no conflict of interest.

## Notes

https://osf.io/ey3ca/

https://osf.io/v9xsw/

https://github.com/corcorana/WIM_HB

### Competing Interest Statement

The authors have declared no competing interest.

### Summary of Updates

Additional analysis of replication dataset collected at the Paris Brain Institute

